# Establishment of a Reverse Genetics System for Influenza D Virus

**DOI:** 10.1101/724732

**Authors:** Hiroho Ishida, Shin Murakami, Haruhiko Kamiki, Hiromichi Matsugo, Akiko Takenaka-Uema, Taisuke Horimoto

**Affiliations:** Department of Veterinary Microbiology, Graduate School of Agricultural and Life Sciences, University of Tokyo, Tokyo, Japan

**Keywords:** bovine respiratory disease complex, influenza D virus, mutant, recombinant virus, reverse genetics, transfection

## Abstract

Influenza D virus (IDV) was initially isolated in the USA in 2011. IDV is distributed worldwide and is one of the causative agents of bovine respiratory disease complex (BRDC), which exhibits high morbidity and mortality in feedlot cattle. Molecular mechanisms of IDV pathogenicity are still unknown. Reverse genetics systems are vital tools not only for studying the biology of viruses, but also for use in applications such as recombinant vaccine viruses. Here, we report the establishment of a plasmid-based reverse genetics system for IDV. We first verified that the 3′-terminal nucleotide of each 7-segmented genomic RNA contained uracil in contrary to the previous report, and were then able to successfully generate recombinant IDV by co-transfecting 7 plasmids containing these genomic RNAs along with 4 plasmids expressing polymerase proteins and NP into HRT-18G cells. The recombinant virus had a growth deficit compared to the wild-type virus, and we determined the reason for this growth difference by examining the genomic RNA content of the viral particles. We found that recombinant virus incorporated an unbalanced ratio of viral RNA segments into particles as compared to the wild-type virus, and thus we adjusted the amount of each plasmid used in transfection to obtain recombinant virus with the same replicative capacity as wild-type virus. Our work here in establishing a reverse genetics system for IDV will have a broad range of applications, including uses in studies focused on better understanding IDV replication and pathogenicity as well as those contributing to the development of BRDC countermeasures.

**IMPORTANCE:** Bovine respiratory disease complex (BRDC) exhibits high mortality and morbidity in cattle, causing economic losses worldwide. Influenza D virus (IDV) is considered to be a causative agent of BRDC. Here, we developed a reverse genetics system that allows for the generation of IDV from cloned cDNAs, and the introduction of mutations into the IDV genome. This reverse genetics system will become a powerful tool for use in studies related to understanding the molecular mechanisms of viral replication and pathogenicity, and will also lead to the development of new countermeasures against BRDC.

## INTRODUCTION

Influenza D virus (IDV), a member of the family *Orthomyxoviridae*, was first isolated from pigs with respiratory illness in Oklahoma, USA, in 2011 (1, 2). Epidemiological analyses revealed that cattle are the main host of the virus, due to their high seroprevalence for IDV (2, 3). Further epidemiological studies revealed that IDVs circulate in cattle in many countries including the USA (2–4), Mexico (5), China (6), Japan (7, 8), France (9), Italy (10), Ireland (11), Luxembourg (12), and African countries (13). Furthermore, serological studies showed that IDV antibodies are found in pigs (14), sheep (15, 16), goats (15, 16), dromedary camels (13, 17), horses (18), and humans (19). These findings imply that IDVs are globally distributed in several animal hosts.

IDV is one of the causative agents of bovine respiratory disease complex (BRDC) (5, 20). BRDC exhibits high morbidity and mortality in feedlot cattle, and over 40% of cattle death in the USA is due to BRDC, causing severe economic losses (21). To control BRDC, combined vaccines consisting of several known causative viral (and bacterial) agents have been employed, but their efficacies are limited (21–23). One possible reason for this reduced efficacy is that the vaccines did not contain all causative agents of BRDC. Recent metagenomic analyses indicate that IDVs are found in BRDC cattle (5). Therefore, addition of IDV to vaccines may help to control BRDC.

Influenza A and B viruses (IAV and IBV) possess 8-segmented negative sense RNA segments (PB2, PB1, PA, HA, NP, NA, M, and NS) as genomes, whereas influenza C virus (ICV) and IDV possess 7-segmented ones (PB2, PB1, P3, HEF, NP, M, and NS) as genomes. The viral RNA (vRNA) of influenza virus forms the ribonucleoprotein complex together with three polymerase subunits, PB2, PB1, and PA/P3, in addition to nucleoprotein (NP). HEF of IDV is a spike protein on the viral envelope that binds the cell surface via terminal 9-O-acetylated sialic acid (24), also known to be a receptor of ICV (25), while the M2 protein of HEF acts as ion channel protein (26). Although the function of M1, NS1, and NS2 proteins of IDV are unknown, we speculate that their functions are similar to those of ICV counterparts (27–30).

Reverse genetics systems are vital tools not only for studying the biology of viruses, but also for use in applications such as recombinant vaccine viruses. While reverse genetics systems for IAV (31–33), IBV (34, 35), and ICV (28, 36) exist, a reverse genetics system for IDV has not been established yet. Such a system could aid in understanding the molecular properties of IDV, in addition to improving methods of disease control. In reverse genetics systems for IAV, IBV, or ICV, RNA polymerase I (PolI) promoter and terminator were used to construct plasmids expressing viral RNAs. These plasmids were co-transfected with plasmids expressing viral-polymerase proteins (PB2, PB1, and PA/P3) and NP into cells, and recombinant viruses were rescued from the supernatants of transfected cells (28, 32, 33, 35, 36). Here, we established a plasmid-based reverse genetics system with IDV using the RNA PolI system, which is similar to that used in reverse genetics systems for other influenza-type viruses.

## RESULTS

### Determination the 3′ terminal sequences of genomic RNA segments of IDV

All influenza viruses possess complementary sequences between the 5′ and 3′ terminal regions of each genome segment. The 5′ and 3′ terminal nucleotides of each segment are adenine (A) and uracil (U), respectively, which are conserved in IAV, IBV, and ICV. However, a previous study has reported that, although 3′ terminal nucleotides of HEF, M, and NS segments of IDV genome are U, those of PB2, PB1, P3, and NP segments are cytosine (C), which are not complementary to A of 5′ terminal nucleotides (1). To confirm these previous findings, we reassessed 3′ and 5′ terminal sequence regions of each segment of D/swine/Oklahoma/1334/2011 (D/OK) through the 3′ or 5′ rapid amplification of cDNA ends (RACE) method. Our results confirmed that the 5′ terminal nucleotides of every segment of IDV were A (Fig. 1A). However, we found that 3′ terminal nucleotides of all segments were U in our D/OK stock, in contrast to those of previous report (1) (Fig. 1B). We next determined 3′ terminal sequences of genome segments from 10 plaque-cloned D/OK and two other IDV stocks of D/bovine/Yamagata/10710/2016 and D/bovine/Nebraska/9-5/2012, and found that segments from all IDVs tested contained U. These data verify that the 3′ terminal sequence of each IDV segment is U, as is seen for other influenza viruses.

**FIG 1.**
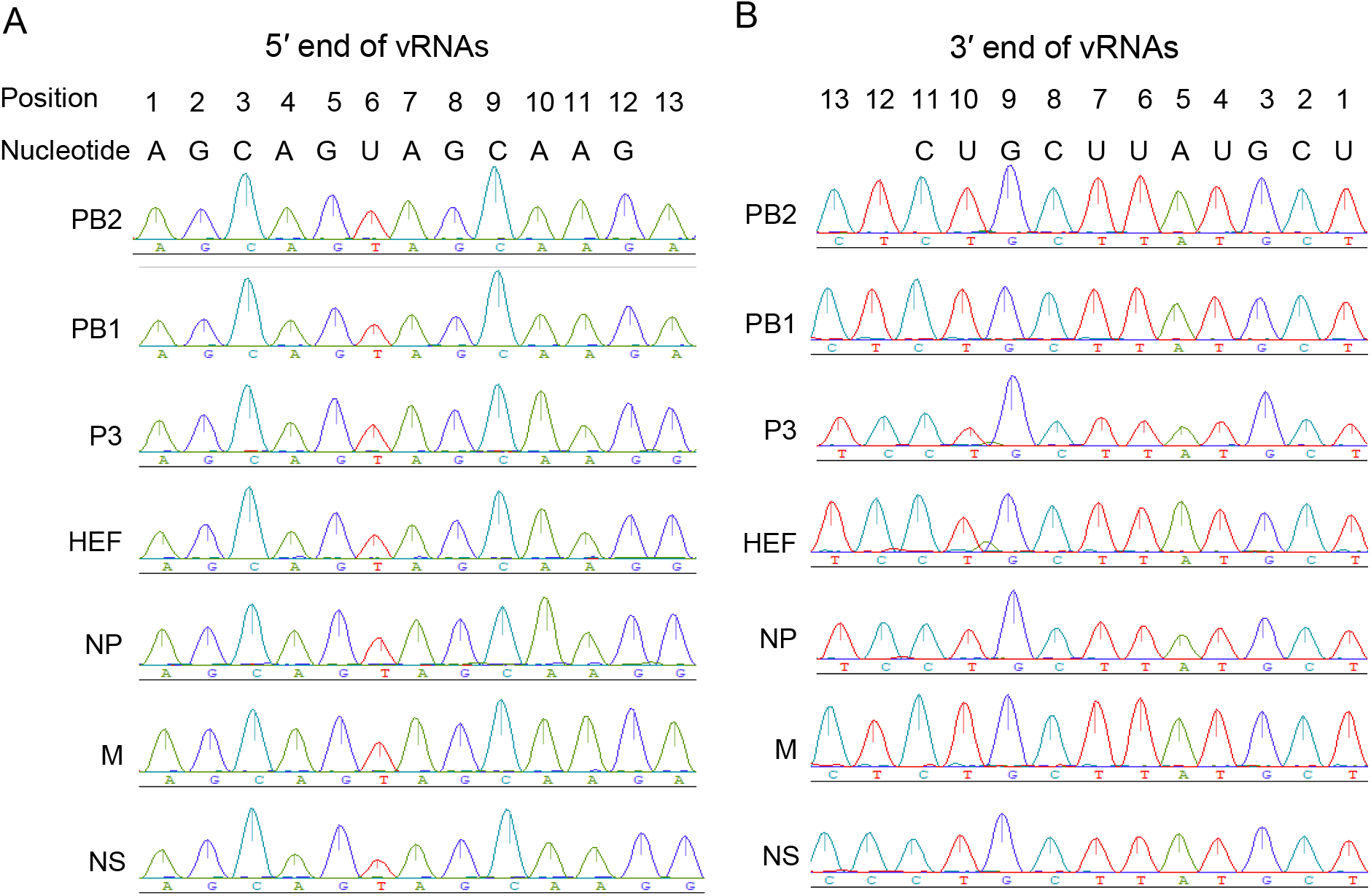
Sequences of 5′ and 3′ terminal regions of viral RNA segments of influenza D virus. The 5′ and 3′ rapid amplification of the cDNA end (RACE) method was performed with D/swine/Oklahoma/1334/2011 (D/OK). (A) Genomic RNA sequences at 5′ terminal regions of RNA segments of D/OK are shown, in which each number indicates the nucleotide position from the 5′ end. (B) Genomic RNA sequences at 3′ terminal regions of RNA segments of D/OK are shown, in which each number indicates the nucleotide position from the 3′ end.

### Generation of recombinant D/OK virus by reverse genetics

Prior to development of a reverse genetics system for IDV, we characterized the D/OK-clone, which was obtained by biological cloning through plaque purification. The growth kinetics and plaque phenotype of the clone were similar to those of the original D/OK stock (Fig. 2). Genome sequences of the D/OK-clone showed two synonymous substitutions in the PB2 and P3 segments, and one nonsynonymous substitution (Ala to Thr) at amino acid position 204 of PB2 when compared to D/OK sequences deposited in the NCBI database (accession number NC_036615-036621) (Table 1).

**FIG 2.**
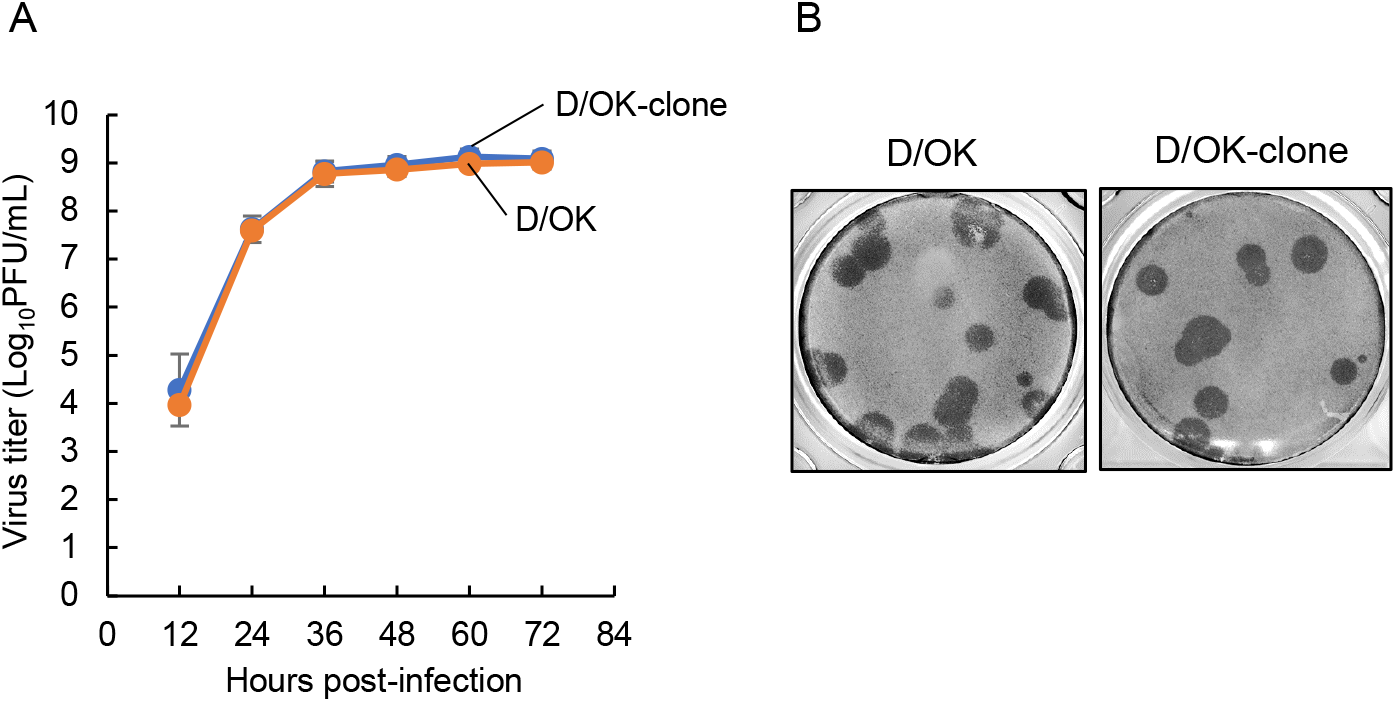
Growth properties of cloned D/OK. (A) Growth kinetics of cloned D/OK (D/OK-clone) were examined in ST cells. D/OK-clone and original D/OK were inoculated onto cells at a multiplicity of infection (MOI) of 0.01. Virus titers were determined at 12 h intervals post-infection by plaque assay, and reported as the mean titer with standard deviations (n=3). (B) Representative plaque morphology of D/OK-clone in ST cells is shown. The plaques were immunologically stained with mouse anti-D/OK polyclonal antibody.

**TABLE 1.** Sequence differences observed between D/swine/Oklahoma/1334/2011 and its plaque-purified virus

| Segment | Amino acid position | Amino acid |  | Nucleotide sequence |  |
| --- | --- | --- | --- | --- | --- |
|  |  | D/OK<sup>a</sup> | D/OK-clone<sup>b</sup> | D/OK | D/OK-clone |
| PB2 | 204 | Ala | Thr | GCA | ACA |
|  | 601 | Ala | Ala | GCG | GCA |
| P3 | 520 | Val | Val | GTC | GTT |

<sup>a</sup> D/OK, wild-type D/swine/Oklahoma/1334/2011
<sup>b</sup> D/OK-clone, a single-clone by three consecutive plaque-purification of D/OK on ST cells.

We cloned cDNAs of 7 vRNA segments of the D/OK-clone into a vRNA synthetic plasmid (referred to as pPolI-D/OK-PB2, -PB1, -P3, -HEF, -NP, -M, and -NS, respectively), and generated 4 vRNA-synthetic plasmids of PB2, PB1, P3, and NP possessing C at their 3′ terminal nucleotides (referred to as pPolI-D/OK-PB2/C, -PB1/C, -P3/C, and -NP/C). We additionally prepared 4 plasmids expressing PB2, PB1, P3, and NP proteins of D/OK (referred to as pCAGGS-PB2, -PB1, -P3, and -NP). We used two sets of vRNA-expression plasmids for reverse genetics; one set consisted of pPolI-D/OK-PB2, -PB1, -P3, -HEF, -NP, -M, and –NS, and the other set consisted of pPolI-D/OK-PB2/C, -PB1/C, -P3/C, -HEF, -NP/C, -M, and -NS. We then transfected HRT-18G cells with 0.2 μg of each plasmid from each set and 4 protein-expression plasmids, pCAGGS-PB2, PB1, P3, and NP. After incubation for 4 days, the supernatant was transferred to fresh swine ST cells and incubated for 4 additional days. Virus rescue was checked by assessing the cytopathic effect (CPE) and performing a hemagglutination (HA) test as well as RT-PCR. We successfully rescued a virus (OK-RG) with the first set of plasmids, but not with the second set, and verified that the OK-RG sequence had no mutations in the genome, including in the 3′ terminal nucleotides of all segments. These data demonstrated the successful generation of recombinant D/OK virus through reverse genetics, but only for instances where the 3′ terminal sequences of vRNAs contained U.

We next compared growth kinetics of the D/OK-clone and OK-RG virus in ST cells (Fig. 3A). Surprisingly, viral titers of OK-RG were 3–4 log lower than those of the D/OK-clone, even though genome sequences of both viruses were identical. Plaque size for the OK-RG virus was smaller than that of the D/OK-clone (Fig. 3B). We passaged OK-RG several times, and purified the virus by plaque picking, but did not see improved growth for OK-RG. This observation suggests the presence of an unknown factor contributing to the inhibition of virus growth in the rescued OK-RG preparation.

**FIG 3.**
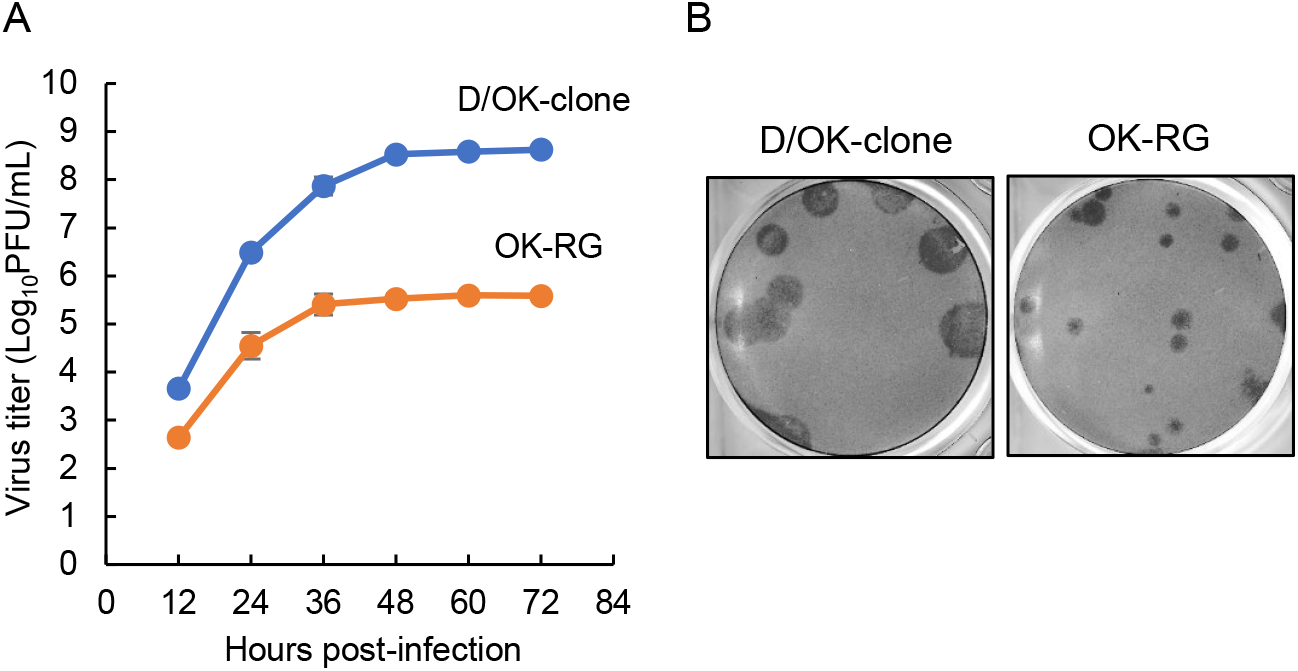
Growth properties of recombinant D/OK. (A) Growth kinetics of recombinant D/OK (OK-RG) generated by reverse genetics were examined in ST cells. OK-RG and D/OK-clone were inoculated onto cells at an MOI of 0.01. Virus titers were determined at 12 h intervals post-infection by plaque assay and reported as the mean titer with standard deviations (n=3). (B) Representative plaque morphology of OK-RG in ST cells is shown. The plaques were immunologically stained with mouse anti-D/OK polyclonal antibody.

### Verification of genomic RNA content packing in OK-RG particles

To gain insight into the mechanism underlying OK-RG growth deficit, we first confirmed vRNA expression in the plasmid-transfected cells. We extracted intracellular vRNAs from the cells and examined content of each RNA segment by northern blot analysis (Fig 4A). vRNA expression varied between segments even though equal amounts of each plasmid were used for transfection, whereas similar amounts of each vRNAs were detected in D/OK clone-infected cells.

**FIG 4.**
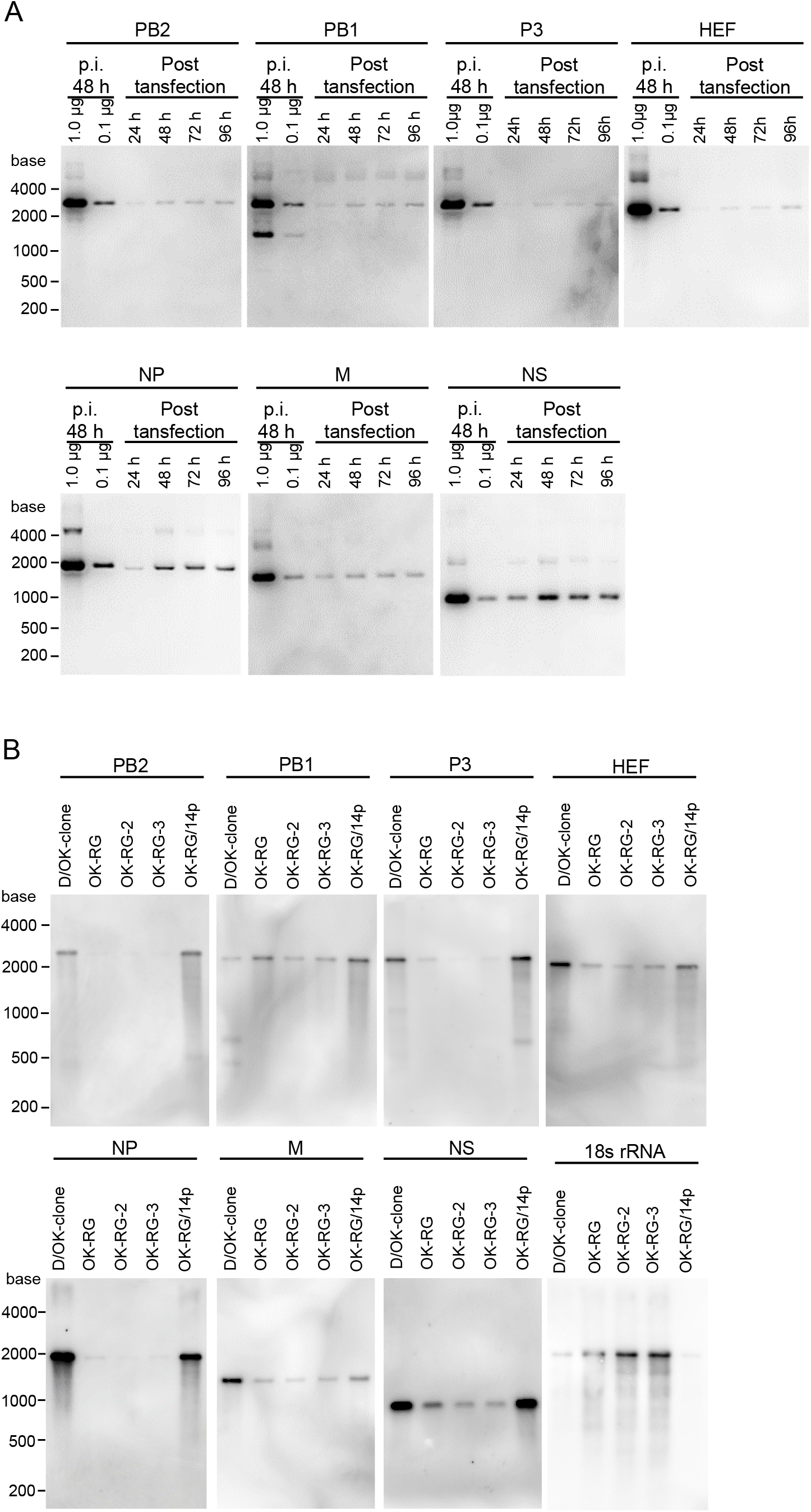
vRNAs in plasmid-transfected cells and in OK-RG particles. (A) Intracellular vRNAs, extracted from plasmid-transfected HRT-18G cells at 24, 48, 72, and 96 h post-transfection, were examined by northern blot analysis with segment-specific riboprobes. vRNAs, extracted from D/OK-clone-infected cells at 48 h post-infection (p.i.), were also examined. (B) RNAs, extracted from the purified D/OK-clone, OK-RG, -RG2, -RG3, and plaque-purified OK-RG (OK-RG/14p), were subjected to northern blot analysis with segment-specific and 18S rRNA-specific riboprobes.

We next examined and compared vRNA content packaged into the D/OK-clone and OK-RGs, including two additional recombinant OK-RG-2 and −3 viruses, which had been rescued independently, by northern blot analysis (Fig. 4B). Interestingly, levels of PB2, P3, and NP RNAs were lower in purified OK-RG particles than in purified D/OK-clone particles, whereas levels of HEF, M, and NS RNAs were slightly lower in OK-RG particles than in D/OK-clone particles. Levels of PB1 RNA were comparable between OK-RG and D/OK-clone particles. These results suggest that vRNA content packaged into OK-RG particles does not reflect vRNA content in the transfected cells.

In our previous report, 7-segmented (HA segment-deficient) IAV incorporated ribosomal RNAs into its virion (37). Thus, we examined the presence of 18S ribosomal RNA (rRNA) in purified recombinant virus, and found a greater amount of 18S rRNAs in OK-RG particles as compared to D/OK-clone particles (Fig. 4B). Taken together, these data demonstrate that OK-RG disproportionally incorporates genomic RNA segments, as well as rRNA, into viral particles. This result suggests that a considerable number of non-infectious particles, such as defective interference particles, are generated in the supernatant through reverse genetics procedures, which may lead to the growth deficit of OK-RG when compared to the D/OK-clone.

To confirm this hypothesis, we directly examined the content of RNA segments packaged into viral particles. We extracted vRNAs from purified OK-RG and separated them through urea-polyacrylamide gel electrophoresis (PAGE), which revealed different profiles of RNA segments for the D/OK-clone and OK-RG (Fig. 5). P3 and NP RNA levels were substantially lower in OK-RG than the D/OK-clone and abundant 18S rRNA, whose band likely covered that of HEF, was detected in OK-RG, as determined through northern blot analysis (Fig. 4B). We found a few extra bands between HEF and NP, and NP and M, suggesting the presence of undefined, deleted forms of RNA segments, which were not detected through northern blot analysis. These data also suggest that OK-RG disproportionally incorporates genomic RNA segments during particle formation.

**FIG 5.**
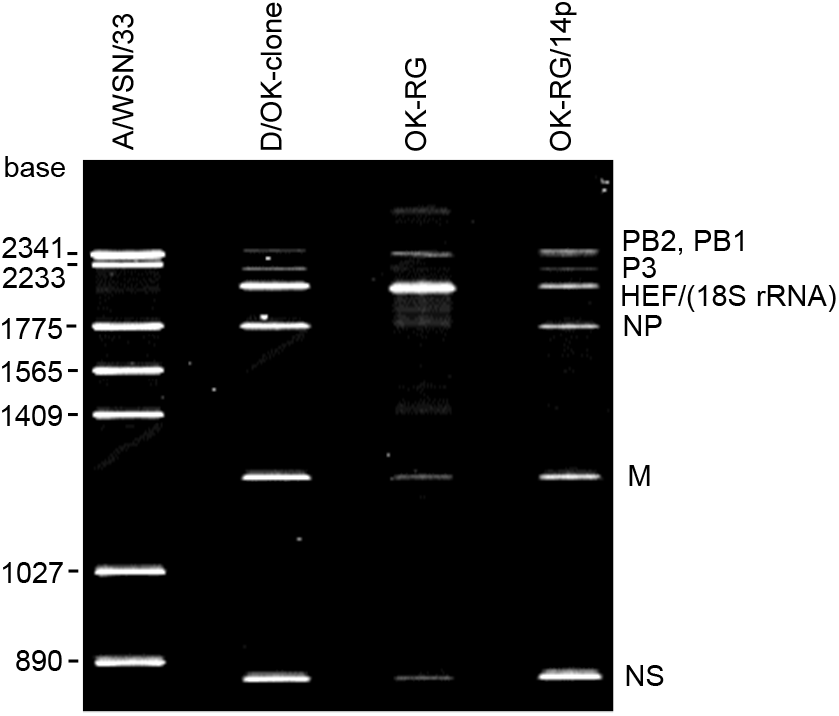
Genomic RNAs in OK-RG particles. RNAs, extracted from the purified D/OK-clone, OK-RG, and OK-RG/14p, were subjected to urea-polyacrylamide gel electrophoresis. Each RNA was run on a 4% urea-polyacrylamide gel for 10 h at 20 mA. The gel was stained with SYBR Gold nucleic acid gel stain.

### Optimization of the reverse genetics system

Based on our northern blot (Fig. 4B) and urea-PAGE (Fig. 5) analyses, we reconsidered transfection conditions for reverse genetics, in order to package similar amounts of each RNA segment into viral particles. We tested an additional 17 transfection conditions by increasing and/or decreasing each vRNA- or protein-expressing plasmids (Table 2). Although viruses were rescued under all conditions, most plaque sizes were smaller than those of the D/OK-clone (Fig. 6A). However, viruses rescued under condition #14 (OK-RG/14) included a high population of those forming large plaques (Fig. 6A). We then plaque-isolated this virus and propagated it once in ST cells (referred to as OK-RG/14p). When growth kinetics of OK-RG/14 and OK-RG/14p were tested in ST cells (Fig. 6B), we found that OK-RG/14p grew at a similar titer to the D/OK-clone, which was approximately 10-fold higher than that of OK-RG/14, suggesting that OK-RG/14p possessed a similar infectivity to wild-type D/OK. In addition, we confirmed that the entire genome sequence of OK-RG/14p was identical to that of the D/OK-clone.

**TABLE 2.**
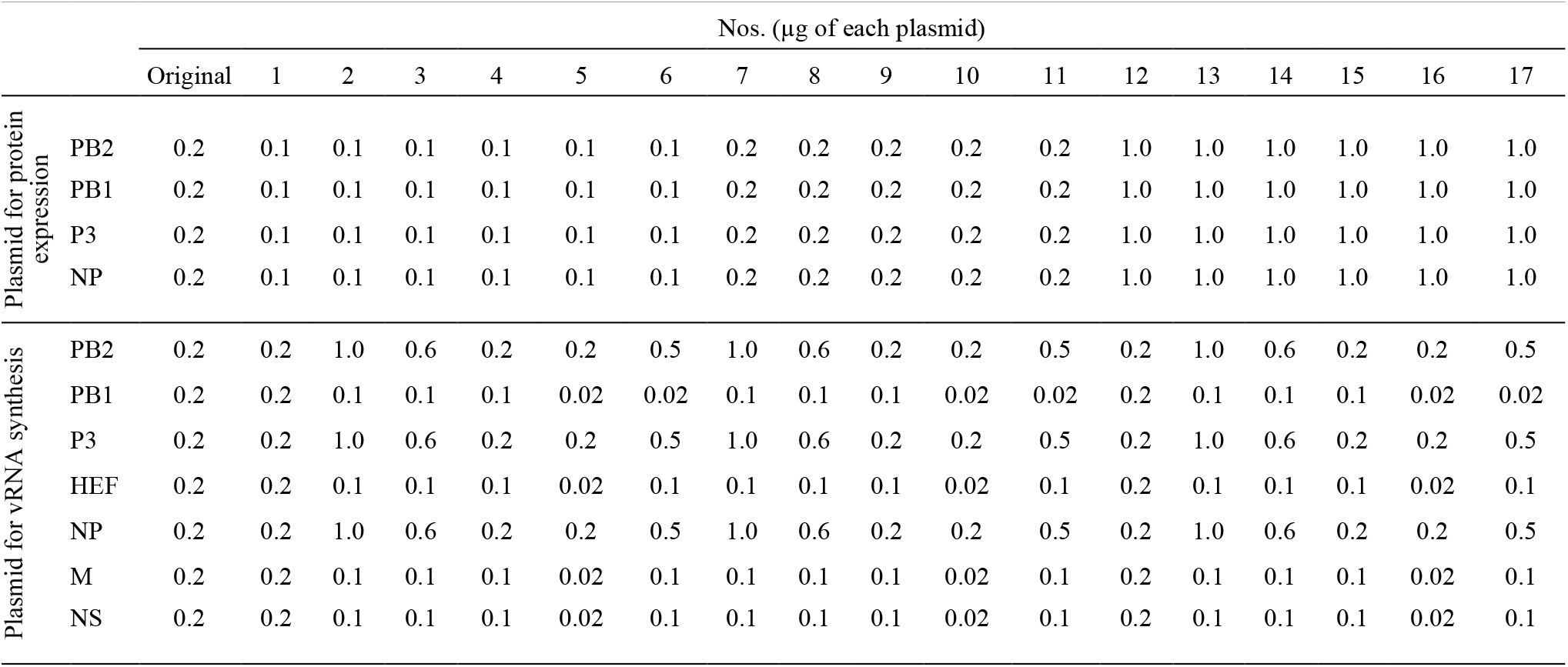
Transfection conditions used for reverse genetics

| | | Nos. ($\mu\text{g}$ of each plasmid) | | | | | | | | | | | | | | | | | |
| --- | --- | --- | --- | --- | --- | --- | --- | --- | --- | --- | --- | --- | --- | --- | --- | --- | --- | --- | --- |
|  |  | Original | 1 | 2 | 3 | 4 | 5 | 6 | 7 | 8 | 9 | 10 | 11 | 12 | 13 | 14 | 15 | 16 | 17 |
| Plasmid for protein expression | PB2 | 0.2 | 0.1 | 0.1 | 0.1 | 0.1 | 0.1 | 0.1 | 0.2 | 0.2 | 0.2 | 0.2 | 0.2 | 1.0 | 1.0 | 1.0 | 1.0 | 1.0 | 1.0 |
|  | PB1 | 0.2 | 0.1 | 0.1 | 0.1 | 0.1 | 0.1 | 0.1 | 0.2 | 0.2 | 0.2 | 0.2 | 0.2 | 1.0 | 1.0 | 1.0 | 1.0 | 1.0 | 1.0 |
|  | P3 | 0.2 | 0.1 | 0.1 | 0.1 | 0.1 | 0.1 | 0.1 | 0.2 | 0.2 | 0.2 | 0.2 | 0.2 | 1.0 | 1.0 | 1.0 | 1.0 | 1.0 | 1.0 |
|  | NP | 0.2 | 0.1 | 0.1 | 0.1 | 0.1 | 0.1 | 0.1 | 0.2 | 0.2 | 0.2 | 0.2 | 0.2 | 1.0 | 1.0 | 1.0 | 1.0 | 1.0 | 1.0 |
| Plasmid for vRNA synthesis | PB2 | 0.2 | 0.2 | 1.0 | 0.6 | 0.2 | 0.2 | 0.5 | 1.0 | 0.6 | 0.2 | 0.2 | 0.5 | 0.2 | 1.0 | 0.6 | 0.2 | 0.2 | 0.5 |
|  | PB1 | 0.2 | 0.2 | 0.1 | 0.1 | 0.1 | 0.02 | 0.02 | 0.1 | 0.1 | 0.1 | 0.02 | 0.02 | 0.2 | 0.1 | 0.1 | 0.1 | 0.02 | 0.02 |
|  | P3 | 0.2 | 0.2 | 1.0 | 0.6 | 0.2 | 0.2 | 0.5 | 1.0 | 0.6 | 0.2 | 0.2 | 0.5 | 0.2 | 1.0 | 0.6 | 0.2 | 0.2 | 0.5 |
|  | HEF | 0.2 | 0.2 | 0.1 | 0.1 | 0.1 | 0.02 | 0.1 | 0.1 | 0.1 | 0.1 | 0.02 | 0.1 | 0.2 | 0.1 | 0.1 | 0.1 | 0.02 | 0.1 |
|  | NP | 0.2 | 0.2 | 1.0 | 0.6 | 0.2 | 0.2 | 0.5 | 1.0 | 0.6 | 0.2 | 0.2 | 0.5 | 0.2 | 1.0 | 0.6 | 0.2 | 0.2 | 0.5 |
|  | M | 0.2 | 0.2 | 0.1 | 0.1 | 0.1 | 0.02 | 0.1 | 0.1 | 0.1 | 0.1 | 0.02 | 0.1 | 0.2 | 0.1 | 0.1 | 0.1 | 0.02 | 0.1 |
|  | NS | 0.2 | 0.2 | 0.1 | 0.1 | 0.1 | 0.02 | 0.1 | 0.1 | 0.1 | 0.1 | 0.02 | 0.1 | 0.2 | 0.1 | 0.1 | 0.1 | 0.02 | 0.1 |

**FIG 6.**
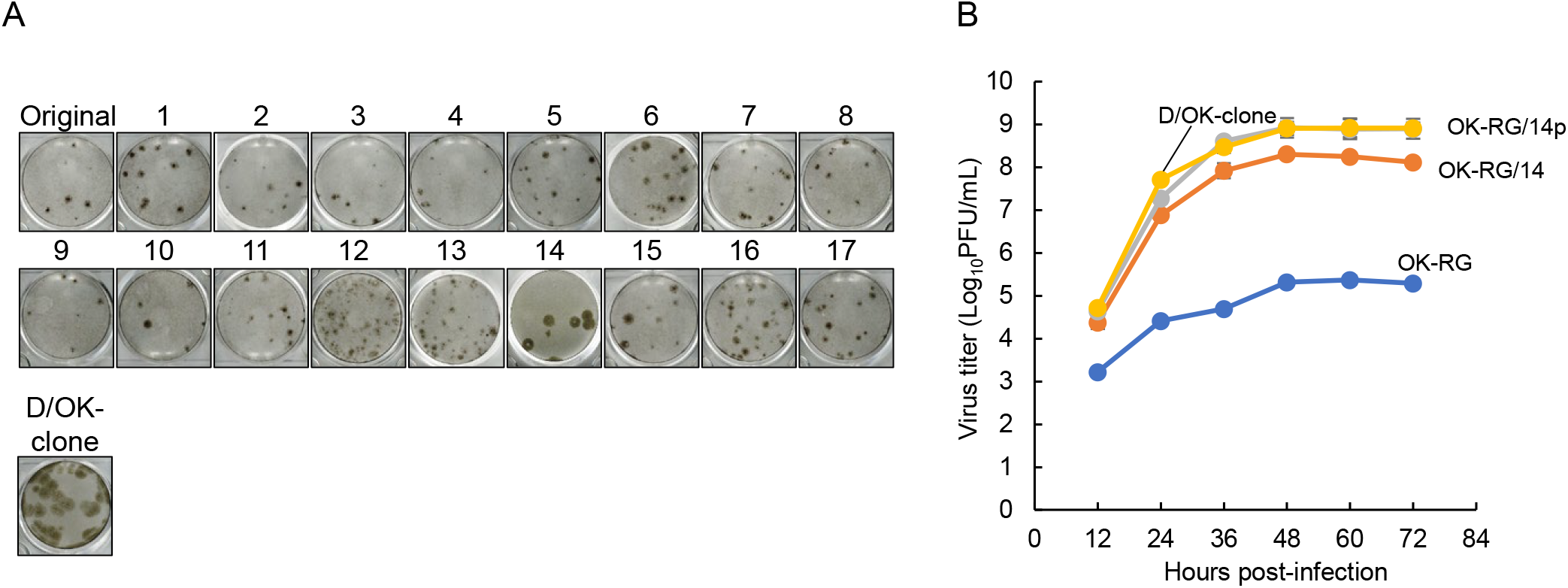
Growth properties of recombinant viruses generated under different conditions in reverse genetics. (A) Representative plaque morphologies of the viruses rescued under both original and 17 different conditions (shown in Table 2), as well as the D/OK-clone, in ST cells are shown. The plaques were immunologically stained with mouse anti-D/OK polyclonal antibody. (B) Growth kinetics of recombinant OK-RG/14 and plaque-purified OK-RG/14p generated under transfection condition #14 were examined. OK-RG/14, OK-RG/14p, OK-RG, and the D/OK-clone were inoculated onto ST cells at an MOI of 0.01. Virus titers were determined at 12 h intervals post-infection by plaque assay, and reported as the mean titer with standard deviation (n=3).

We next assessed content of RNA segments packaged into purified OK-RG/14p particles. In a northern blot analysis, the RNA segment content of OK-RG/14p was similar to that of the D/OK-clone, and 18S rRNA was not detected in OK-RG/14p particles (Fig. 4B). In a urea-PAGE analysis, the profiles of RNA segments were similar between D/OK-clone and OK-RG/14p, except for a greater amount of PB2 and/or PB1 RNA packaging in OK-RG/14p particles (Fig. 5). These results suggest that a well-balanced packaging of RNA segments can be achieved under transfection condition #14, leading to similar growth properties for OK-RG/14p and the D/OK-clone.

## DISCUSSION

Reverse genetics systems, which are used to generate recombinant viruses, are vital tools not only to study the biology of influenza viruses but also for antiviral strategies such as generation of recombinant vaccine viruses. Here, we established a plasmid-based reverse genetics system for IDV.

For previously published reverse genetics systems with IAV, IBV, and ICV, human embryonic kidney (HEK) 293T cells were mainly used for virus rescue due to their high transfection efficiency and presence of T antigen required for replication of pCAGGS-based vectors (28, 32, 33, 35, 36). In this study, we first tested this cell line for potential use in an IDV reverse genetics system. Although recombinant D/OKs were rescued after several attempts, their infectivities (PFU titers) were low, despite high HA titers in the supernatant of transfected cells. Their plaque sizes were also small, suggesting the presence of abundant non-infectious particles. Therefore, we next tested HRT-18G cells, which have been widely used for IDV isolation, in our IDV reverse genetics system. Although this cell line showed a much lower transfection efficiency than HEK293T cells, recombinant D/OKs with high infectivity were rescued, as revealed by higher PFU/HA ratios than those seen with HEK293T cells. These findings imply that support of viral replication is a more important factor to consider than transfection efficiency when choosing cells for use in an IDV reverse genetics system. Therefore, a possible system in which the bovine (or swine) PolI vector is utilized as a vRNA-synthetic plasmid for transfection in bovine (or swine) cells may enhance rescue efficiency of recombinant IDVs.

IAV, IBV, and ICV contain adenine (A) and uracil (U) at the 5′ and 3′ terminal ends of every RNA segment, respectively, and complementary sequences of both terminal regions, including these two nucleotides, form panhandle structures of viral ribonucleoprotein (RNP), as well as transcription/replication promoter structures for the viral genome in IAV (38). Here, we found that all RNA segments of IDV also possess U at their 3′ terminal ends, in contrast to previous report showing that PB2, PB1, P3, and NP RNAs of D/OK possessed cytosine (C) (1). Although we were not able to rescue recombinant D/OK with RNA segments possessing C at the 3′ terminal ends, this does not eliminate the possibility that the original D/OK may possess C at the 3′ terminal ends of these RNA segments. However, it is likely that viral passages in laboratory would result in selection of D/OK with U at the 3′ terminal ends of all RNA segments. Alternatively, C at the 3′ terminal end may be incorrectly read due to an undefined artifact resulting from the RACE method (1). Further sequence analyses of the 3′ terminal ends of RNA segments in other multiple IDV isolates will clarify this point.

In this study, we successfully established a reverse genetics system for IDV by fine adjustment of quantitative transfection conditions for viral RNA-synthetic and protein-expression plasmids. This adjustment resulted in balanced packaging of 7 RNA segments in viral particles, leading to the rescue of recombinant D/OK with growth properties similar to the D/OK-clone. This finding suggests a different mechanism for genome packaging between IAV and IDV. Two models for genome packaging of influenza virions have been proposed: random and selective packaging (39). Previous studies support that IAV most likely contains a selective packaging mechanism (39–42). IAV and IBV incorporate eight RNPs arranged in a specific pattern “1+7”, in which seven RNPs surround the central one (40, 43, 44). Surprisingly, artificially generated 7-segmented IAV, which lacks an HA segment, incorporated rRNA as the eighth RNP instead of an HA segment (37). Interestingly, ICV and IDV both possess a 7-segmented genome, yet also incorporated 8 RNPs, although inclusion of rRNA in virions was not assessed (45). These findings imply that all influenza viruses may incorporate 8 RNPs, suggesting the presence of an undefined random packaging mechanism in addition to a selective packaging one, especially for ICV and IDV. Several studies also support the that the genome packaging mechanisms of IAV and ICV are different. In one study, artificially generated virus-like particles (VLPs) of IAV with less than 8 RNA segments were less efficiently produced than those with one complete set of 8 RNA segments (46, 47), whereas in a different study, VLPs of ICV with single RNA segments were produced more efficiently than those with a complete set of 7 RNA segments (27, 48). Our present study also supports such a difference, in which OK-RG incorporated unbalanced ratios of RNA segments and rRNA into particles when equal amounts of plasmids were used for transfection. This imbalance may have resulted in the generation of abundant non-infectious particles in the supernatant of transfected cells, which may have interfered with virion production. However, this transfection condition is regularly used for reverse genetics with IAV, suggesting a difference in genome packaging between IAV and IDV. Further studies are required to elucidate differences in the genome packaging mechanism between 8-segmented and 7-segmented viruses.

In conclusion, this study established a reverse genetics system for IDV. However, transfection procedures must be carefully observed, since we determined that only one transfection condition (#14) can efficiently generate virions with original infectivity. Nonetheless, this system will allow us to produce recombinant viruses with viable mutation(s) in the IDV genome, and will aid in the study of molecular mechanisms of virus replication, including genome packaging and pathogenicity. Based on such knowledge, we will be able to generate live attenuated IDVs, which could be an effective control measure against BRDC.

## MATERIALS AND METHODS

### Cells and viruses

Human embryonic kidney 293T cells (obtained from Riken BRC, RCB2202), human rectal tumor HRT-18G cells (obtained from ATCC, CRL-11663), and swine testis (ST) cells (obtained from ATCC, CRL-1746) were maintained in Dulbecco’s Modified Eagle’s Medium (DMEM; Fujifilm Wako Pure Chemical, Osaka, Japan) supplemented with 10% fetal bovine serum (FBS) at 37°C. D/swine/Oklahoma/1334/2011 (D/OK) (1) and D/bovine/Nebraska/9-5/2012 (4) were kindly provided by Dr. B. Hause (Kansas State University). Our isolated D/bovine/Yamagata/10710/2016 (45) was also used. These IDVs were propagated in ST cells in Eagle’s Minimum Essential Medium (MEM; Life Technologies/Gibco, Paisley, UK) containing 0.3% bovine serum albumin (MEM/BSA) supplemented with 0.5 µg/ml L-1-tosylamido-2-phenyl chloromethyl ketone (TPCK)-trypsin (Worthington, Lakewood, NJ, USA) and stocked at −80°C.

### Construction of plasmids for reverse genetics

RNA was extracted from culture supernatant of plaque-cloned D/OK (D/OK-clone)-infected ST cells using ISOGEN-LS (Nippon Gene, Tokyo, Japan). The extracted RNA was reverse-transcribed using PrimeScript reverse transcriptase (Takara Bio, Shiga, Japan), and the cDNAs were amplified by PCR using KOD FX Neo (Toyobo, Osaka, Japan) and a set of segment-specific primers containing 15 nucleotide-overlapped sequences to the cloning site of pHH21 plasmid, which contains human RNA polymerase I (PolI) promoter and murine PolI terminator sequences (33). The PCR products were purified by a FastGene Gel/PCR extraction kit (Nippon Genetics, Tokyo, Japan) and cloned into *Bsm*BI-digested pHH21 using a Gibson assembly master mix (New England BioLabs (NEB) Japan, Tokyo, Japan), resulting in pPol-D/OK-PB2, -PB1, -P3, -HEF, -NP, -M, and -NS plasmids. pPolI-D/OK-PB2/C, -PB1/C, -P3/C, and -NP/C plasmids possessing C at the 3′ terminal ends of viral RNAs were also generated using a set of segment-specific primers containing C at the 3′ terminal end, and 15 nucleotide-overlapped sequences to the cloning site of pHH21. Coding regions of PB2, PB1, P3, and NP segments were PCR-amplified using KOD FX Neo and 15 nucleotide-overlapped sequences to the cloning site of pCAGGS plasmid (49). The PCR products were cloned into the *Eco*RI and *Xho*I-digested pCAGGS using a Gibson assembly master mix, resulting in pCAGGS-D/OK-PB2, -PB1, -P3, and -NP plasmids.

### Generation of recombinant IDV by reverse genetics

Sixty percent confluent HRT-18G cells, seeded on a 6-well plate, were transfected with 7 viral RNA-synthetic plasmids and 4 protein (PB2, PB1, P3, and NP)-expression plasmids using 0.2 µg of each plasmid with 11 µl of 1 µg/µl PEI MAX (Polysciences, Warrington, PA, USA). Prior to transfection, the plasmid DNAs and transfection reagent were mixed and incubated at 23°C for 20 min. The mixtures were then added to the cells and incubated at 37°C. At 2 days post-transfection, the supernatants were removed, and cells were washed twice with MEM before addition of 2 mL of MEM/BSA containing 0.5 µg/ml TPCK-trypsin. After 3 days of incubation at 37°C, the supernatants were collected, diluted 10-fold with MEM/BSA, and inoculated on ST cells for 1 h. The cells were washed twice in MEM before addition of 2 mL of MEM/BSA containing 0.5 µg/ml TPCK-trypsin, and incubated at 37°C. At 2–3 days post-infection, supernatants were collected, and viruses were titrated by a plaque assay in ST cells.

### Plaque assay

Confluent ST cells on a 12-well plate were washed twice with phosphate-buffered saline (PBS), inoculated with 0.1 mL each of serial 10-fold diluted viruses in MEM/BSA, and incubated for 1 h at 37°C. Cell were then washed with MEM/BSA, covered with 1 mL of MEM/BSA containing 1% Seakem GTG agarose (Lonza Japan, Chiba, Japan) and 0.5 µg/ml TPCK-trypsin, and incubated at 37°C for 3 days. 0.5 mL of 30% formalin in PBS was added to each well for fixation at 4°C overnight. After formalin and agarose were removed, the cells were washed with PBS and permeabilized with 0.1% Triton X-100 in PBS for 15 min at 23°C. After blocking with Block Ace (KAC, Hyogo, Japan), the cells were incubated with anti-IDV mouse immune sera as a primary antibody for 60 min, followed by incubation with biotinylated anti-mouse IgG antibody (#B7264; Sigma, Kanagawa, Japan) for 30 min, followed by a complex containing 4 µg/ml biotinylated peroxidase (Invitrogen/Thermo Fisher Scientific, Tokyo, Japan) and 8 µg/ml streptavidin (Fujifilm Wako Chemicals) for 30 min. The plaques were visualized by the DAB peroxidase substrate kit (Vector Laboratories, Burlingame, CA, USA), according to the manufacturer’s instruction.

### Hemagglutination assay

A hemagglutination (HA) assay was performed in U-bottom 96-well microplates as previously described (50). Briefly, serial 2-fold dilutions of the supernatants in 50 µL of PBS were mixed with 50 µL of 0.7% turkey red blood cells and incubated for 30 min at 23°C before reading. The HA titers were determined as the reciprocal of the highest virus dilution showing complete HA.

### 5′ and 3′ RACE

5′ and 3′ rapid amplification of the cDNA end (RACE) was performed using a 5′/3′ RACE kit (2nd Generation; Roche, Basel, Switzerland) with some modifications. Briefly, viral RNAs were extracted from purified virus particles using ISOGEN-LS. For 5′ RACE, the extracted RNAs were reverse-transcribed using segment-specific primers. The cDNA products were then added along with poly-dA using terminal deoxynucleotidyl transferase (Roche). The poly-dA-added cDNAs were amplified by segment-specific primers and our oligo-dT primer containing a synthetic tag (5′-GACCACGCGTATCGATGTCGACTTTTTTTTTTTTTTTT-3′) (instead of oligo d(T)-anchor primer (5′-GACCACGCGTATCGATGTCGACTTTTTTTTTTTTTTTTV-3′) provided by the kit), followed by semi-nested-PCR using segment-specific primers and PCR anchor primer (5′-GACCACGCGTATCGATGTCGAC-3′). For 3′ RACE, a poly-A tail was added to the viral RNAs, using *E. coli* poly A polymerase (NEB). The poly-A-added RNAs were reverse-transcribed using the tag-added oligo-dT primer and the cDNAs were amplified by PCR using PCR anchor primer and segment-specific primers. The PCR products were sequenced using a 3130xl Genetic Analyzer (Life Technologies Japan/Applied Biosystems, Tokyo, Japan). The sequences of the segment-specific primers will be provided upon request.

### Virus purification

The virus-containing supernatants from ST cells were clarified by low-speed centrifugation to remove cell debris, and concentrated by ultracentrifugation at 40,000 ×g for 2 h at 4°C using a P19A rotor (Himac, Tokyo, Japan). The concentrated viruses were resuspended in PBS and layered onto a 20%, 30%, 50%, and 60% discontinuous sucrose gradient and ultracentrifuged at 110,000 ×g for 3 h at 4°C using a P32ST rotor (Himac). The virus-containing interface between the 30% and 50% sucrose gradient was collected, diluted in PBS, and ultracentrifuged once more at 110,000 × g for 3 h at 4°C using a P32ST rotor. Purified virus pellets were suspended in a small amount of PBS.

### Urea-PAGE

RNAs of the purified viruses were extracted using an miRNeasy mini kit (Qiagen, Tokyo, Japan). To separate RNA segments, urea-PAGE was used as previously described (45) with some modifications. One hundred ng of viral RNAs were run on a 4% polyacrylamide gel containing 7 M urea. The gel was then stained with SYBR Gold nucleic acid gel stain (Invitrogen). The gel images were obtained by a gel imager (Printgraph; ATTO, Tokyo, Japan).

### Northern blot analysis

Northern blot analysis was performed as previously described (37) with some modifications. Briefly, viral RNAs were extracted from purified viruses, plasmid-transfected, or virus-infected HRT-18G cells using ISOGEN reagents (Nippon Gene). Viral (10 ng) or intracellular (1 µg) RNAs were denatured in denaturing buffer (50% formamide and 17.8% formalin in 1% MOPS buffer), separated on 1.5% denaturing agarose-formaldehyde gels, and transferred onto a nylon membrane with a molecular weight marker (BioDynamics Laboratory, Tokyo, Japan). Biotin-labeled strand-specific RNA probes for PB2 (hybridizing to nucleotides (nt) 15 to 1257), PB1 (nt 26 to 1260), P3 (nt 23 to 1038), HEF (nt 27 to 1017), NP (nt 28 to 954), M (nt 29 to 1220), NS (to nt 29 to 868) vRNA, and 18S rRNA were synthesized using a biotin-11-UTP (Roche) and a T7 RNA expression kit (Promega, Madison, WI, USA), according to the manufacturer’s instructions. Detection of RNA segments was performed using an ABC kit (Vector Laboratories) and a DIG block and wash buffer set (Roche), according to the manufacturer’s instructions. The signals were detected with Clarity ECL substrate (Bio-Rad, Hercules, CA, USA), and blot images were obtained by Image Quant LAS 4000mini system (Fujifilm, Tokyo, Japan).

## ACKNOWLEDGMENTS

We thank Dr. Ben M. Hause at Kansas State University for providing viruses and Dr. Yoshihiro Kawaoka at University of Tokyo for providing plasmids. S.M. received a Grant-in-Aid for Encouragement of Young Scientists (A) (grant number: 17H05042) by Grants-in-Aid from the Japan Society for the Promotion of Science. T.H. received, in part, by a Grant-in-Aid for Scientific Research (A) (grant number: 18H03971) by Grants-in-Aid from the Japan Society for the Promotion of Science, and by Livestock Promotion Funds from the Japan Racing Association.

